# The *white* gene controls copulation success in *Drosophila melanogaster*

**DOI:** 10.1101/072710

**Authors:** Chengfeng Xiao, Shuang Qiu, R Meldrum Robertson

## Abstract

Characteristics of male courtship behavior in *Drosophila melanogaster* have been well-described, but the genetic basis of male-female copulation is largely unknown. Here we show that the *white* (*w*) gene, a classical gene for eye color, is associated with copulation success. 82.5% of wild-type Canton-S flies copulated within 60 minutes in circular arenas, whereas few white-eyed mutants mated successfully. The *w*^+^ allele exchanged to the X chromosome or duplicated to the Y chromosome in the white-eyed genetic background rescued the defect of copulation success. The *w*^+^-associated copulation success was independent of eye color phenotype. Addition of the mini-*white* (m*w*^+^) gene to the white-eyed mutant rescued the defect of copulation success in a manner that was m*w*^+^ copy number-dependent. Lastly, male-female sexual experience mimicked the effects of *w*^+^/m*w*^+^ in improving successful copulation. These data suggest that the *w*^+^ gene controls copulation success in *Drosophila melanogaster*.

## Introduction

Mating behavior in wild-type *Drosophila* consists of a series of courtship rituals and subsequent copulation. Successful copulation is a consequence of sexual interactions between specific male stimuli and appropriate female responses^1^. Mating success is of great importance to the fitness of a population. Behavioral features of mating in *Drosophila* are well-described^2, 3^. Many of the behavioral components are quantifiable and have been genetically attributed to chromosomal arrangement or polymorphism^4–8^, and even individual genes^9–15^. For example, in *Drosophila persimilis* and *Drosophila pseudoobscura*, males carrying the commonest gene arrangement on the third chromosome mate rapidly, whereas males with the less frequent arrangement mate slowly^5, 6^.

The *white* (*w*^+^) gene in *Drosophila*, discovered in 1910 by Thomas Hunt Morgan^16^, encodes a subunit of an ATP-binding cassette (ABC) transporter, which loads up pigment granules and deposits the content to pigment cells in the compound eyes, ocelli, Malpighian tubules and testis^17, 18^. In addition, the White protein transports bioamines, neurotransmitters, metabolic intermediates, second messengers and many small molecules^15, 19–23^. White is thus proposed to have housekeeping functions in the central nervous system besides its classical role in eye pigmentation^15, 21, 24, 25^.

It was not long after the discovery that the mutant form of *w*^+^ was found to be involved in sexual discrimination. Sturtevant (1915) reported that Morgan’s white-eyed male flies^16^ were less successful than wild-type males in mating females^2^. A ratio of copulation success is 0.75 for white-eyed male to 1 for wild-type male^26^. Such a sexual discrimination against white-eyed males eventually results in the elimination of mutant allele from a laboratory population^26^. Although these findings suggest a role for *w* in mating selection, there is no direct evidence whether *w*^+^ determines successful male-female copulation.

Ectopic expression or intracellular mislocation of White protein, a product of mini-*white* (m*w*^+^), induces male-male courtship chaining^12, 13, 15^. However, m*w*^+^ males do not reduce courtship preference for females^15, 27^. Also, there are no data whether m*w*^+^ has an effect on copulation success. A complete lack of White reduces sexual arousal of males in the daylight but not in the dark^28^, but whether or not the reduced sexual arousal affects copulation success, more specifically, whether *w*^+^ or m*w*^+^ promotes successful male-female copulation is still unknown.

In the current study we examined copulation success in wild-type Canton-S (CS) and white-eyed mutant (*w*^1118^) strains. We demonstrate that *w*^1118^ flies have a defect of copulation success in small circular arenas. The defect can be rescued by several genetic approaches, including the exchange of *w*^+^ allele from wild-type to *w*^1118^ background; *w*^+^ duplication to the Y chromosome of *w*^1118^ flies; and transgenic insertion of a m*w*^+^ to the *w*^1118^ background. We further show that homozygous m*w*^+^ alleles over-rectify the reduced courtship activities of *w*^1118^ males, and that there is a positive correlation between m*w*^+^ copies and copulation success in flies with a *w*^1118^ genetic background.

## Results

### A defect of copulation success in *w* mutants

When a naive CS male (4-day-old) and a virgin CS female (4-day-old) were placed into a circular arena (1.27 cm diameter and 0.3 cm depth), they initiated courtship activities and started copulation (flies in copulo) within minutes. Such a sexual interaction was highly consistent from one pair to another (Fig. 1A). In the *w* mutant *w*^1118^, however, a naive male and a virgin female together rarely initiated typical courtship activities and failed to copulate within 60 min. The failure of copulation in *w*^1118^ was consistent between pairs (Fig. 1B). The difference of copulation success was clearly observable between two strains. We thus explored the contribution of *w*^+^ to the copulation success in *Drosophila*.

**Figure 1.**
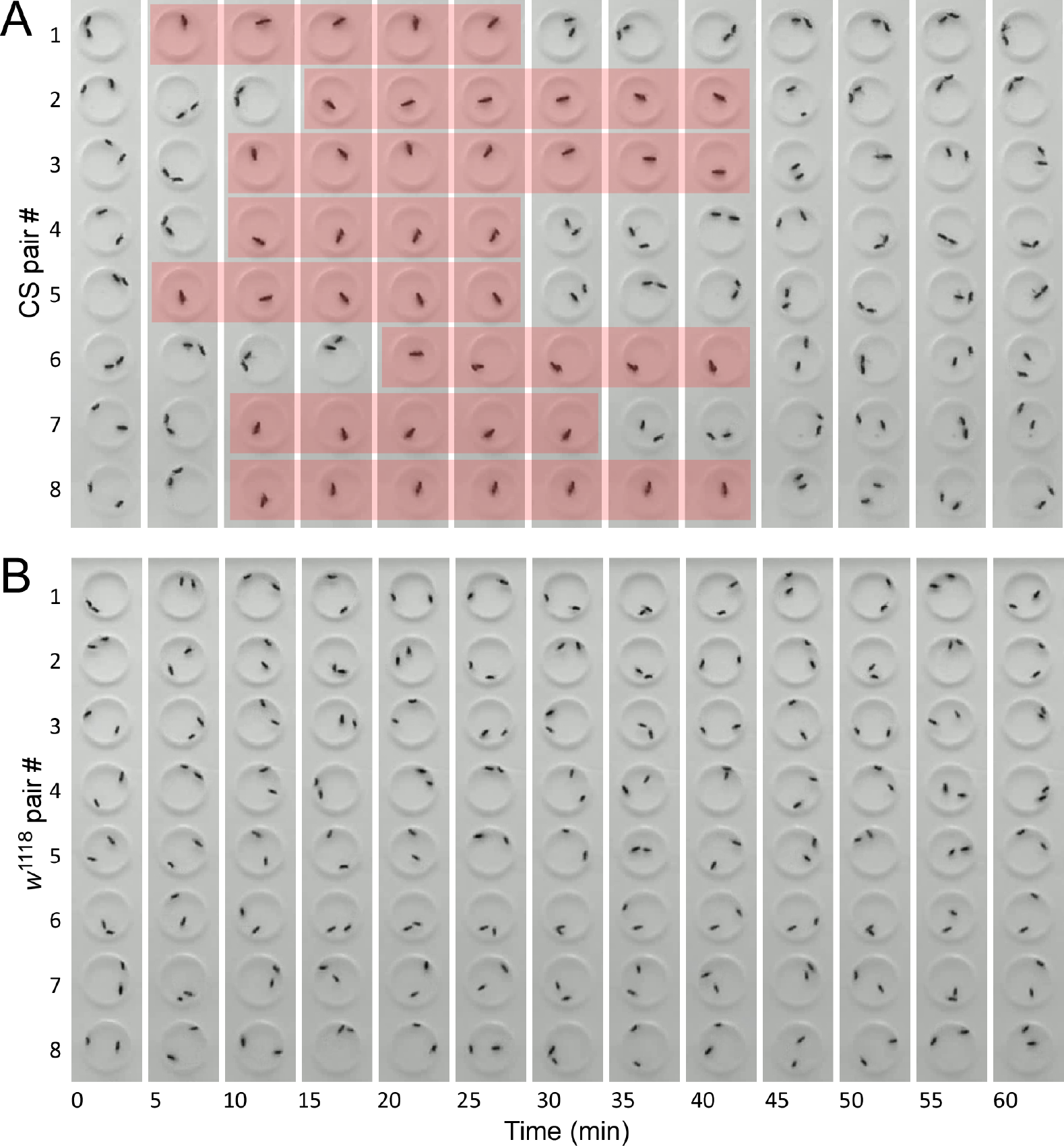
A defect of copulation success in *w*^1118^ flies. Shown are the video-frames sampled once per five minutes. (**A**) Copulation success of wild-type (CS) in the circular arenas. Each pair of flies (a naive male and a virgin female) were loaded into a circular arena (1.27 cm diameter, 0.3 cm depth), and their copulation activities within 60 minutes were examined. Successful copulation is highlighted with light-red. Tested flies are at 4-7 days old. (**B**) A defect of copulation success in *w*^1118^ flies. Naive males and virgin females of *w*^1118^ flies were used.

In CS flies, the percentage of copulation success was 82.5% (33/40) (Fig. 2A and 2E). Median copulation duration was 25.4 min (interquartile range (IQR) 22.3 - 28.6 min) with a median latency of 11.1 min (IQR 6.6 - 19.3 min) (Fig. 2A). In *w*^1118^ flies, copulation success within 60 min was 0% (0/40) (Fig. 2B and 2E), a level significantly lower than that in CS (*P <* 0.0001, Fisher’s exact test). Copulation success of CS flies in the circular arenas was comparable with previous observations that several wild-types had successful copulation rates of 50-90% within 60 min in varying sized mating chambers^29–32^. *w*^1118^ flies displayed a defect of copulation success in the circular arenas within 60 min.

**Figure 2.**
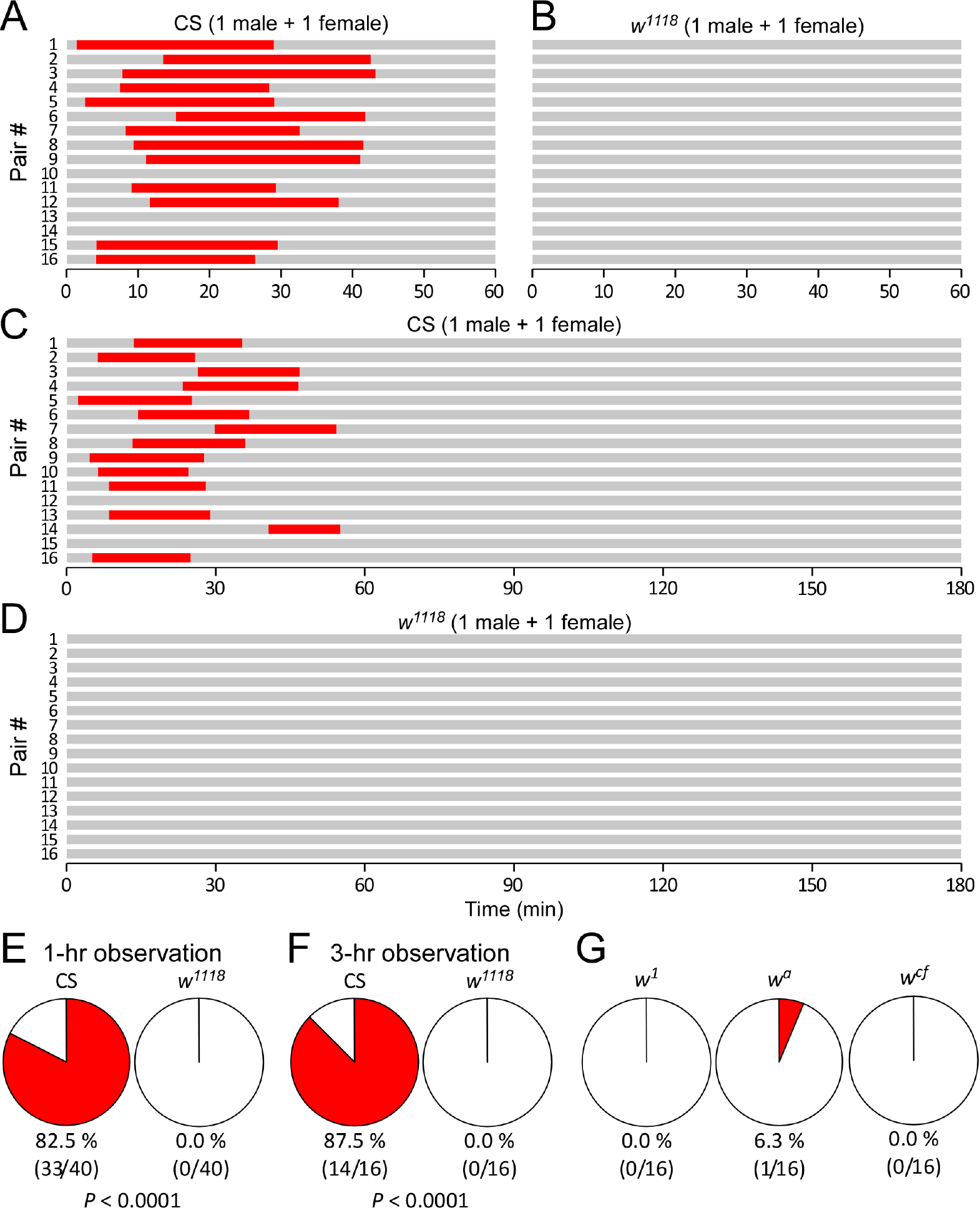
Reduced copulation success in *w* mutants. **(A, B)** Schematic illustrations of copulation success within 60 min in CS and *w*^1118^ flies. The time for successful copulation (red) and non-copulation (grey) are indicated. (**C, D**) Copulation success within 180 minutes in CS and *w*^1118^ flies. (**E**) Percentages of copulation success (red) during 1-hr observation in CS and *w*^1118^ flies. (**F**) Percentages of copulation success during 3-hr observation in CS and *w*^1118^ flies. (**G**) Percentages of copulation success in *w*^1^, *w^a^* and *w^c f^* mutants. Notes: The red piece in the pie chart indicates copulation success. A pair of flies comprise a naive male and a virgin female. Tested flies are at 4-7 days old. Numbers of pairs tested and pairs with successful copulation are shown. *P* values are from Fisher’s exact tests.

CS pairs, if successful in copulation, finished within 60 min. It is possible that *w*^1118^ flies require longer than an hour to start copulation. We then examined the copulation success of flies in the circular arenas for 180 min. CS pairs copulated once with a successful rate of 87.5% (14/16), which was comparable with the successful rate observed during 60-min experiments (*P >* 0.05, Fisher’s exact test). CS flies, once again if successful, finished copulation within the first 60 min. There was no second copulation during the next 120 min (Fig. 2C and 2F). This was consistent with previous reports^2, 30, 33^, and proved the sufficiency of 60-min observation for copulation success in wild-type. As a contrast, *w*^1118^ pairs showed a complete lack of copulation within 180 min (Fig. 2D and 2F). Therefore, the defect of copulation success in *w*^1118^ flies was further evident within a period of prolonged observation.

The defect of copulation success was also observed in several additional *w* mutants, including *w*^1^ (0.0%, 0/16) (*P <* 0.0001 compared with CS, Fisher’s exact test), *w^a^* (6.3%, 1/16) (*P <* 0.0001 compared with CS, Fisher’s exact test) and *w^c f^* (0.0%, 0/16) (*P <* 0.0001 compared with CS, Fisher’s exact test) (Fig. 2G). Taken together, *w* mutants displayed a defect of copulation success in the circular arenas.

### *w*^1118^ male showed a severe defect of copulation success

We next examined which sex of *w*^1118^ flies contributed largely to the defect of copulation success. The rate of successful copulation between CS male and *w*^1118^ female (60.0%, 6/10) was comparable to that between CS male and CS female (82.5%, 33/40) with no significant difference (*P* = 0.197, Fisher’s exact test) (Fig. 3A). However, copulation between *w*^1118^ male and CS female (0.0%, 0/10), and that between *w*^1118^ male and *w*^1118^ female (0.0%, 0/40) were both unsuccessful. Thus *w*^1118^ male but not female displayed a severe defect for copulation success in the circular arena.

**Figure 3.**
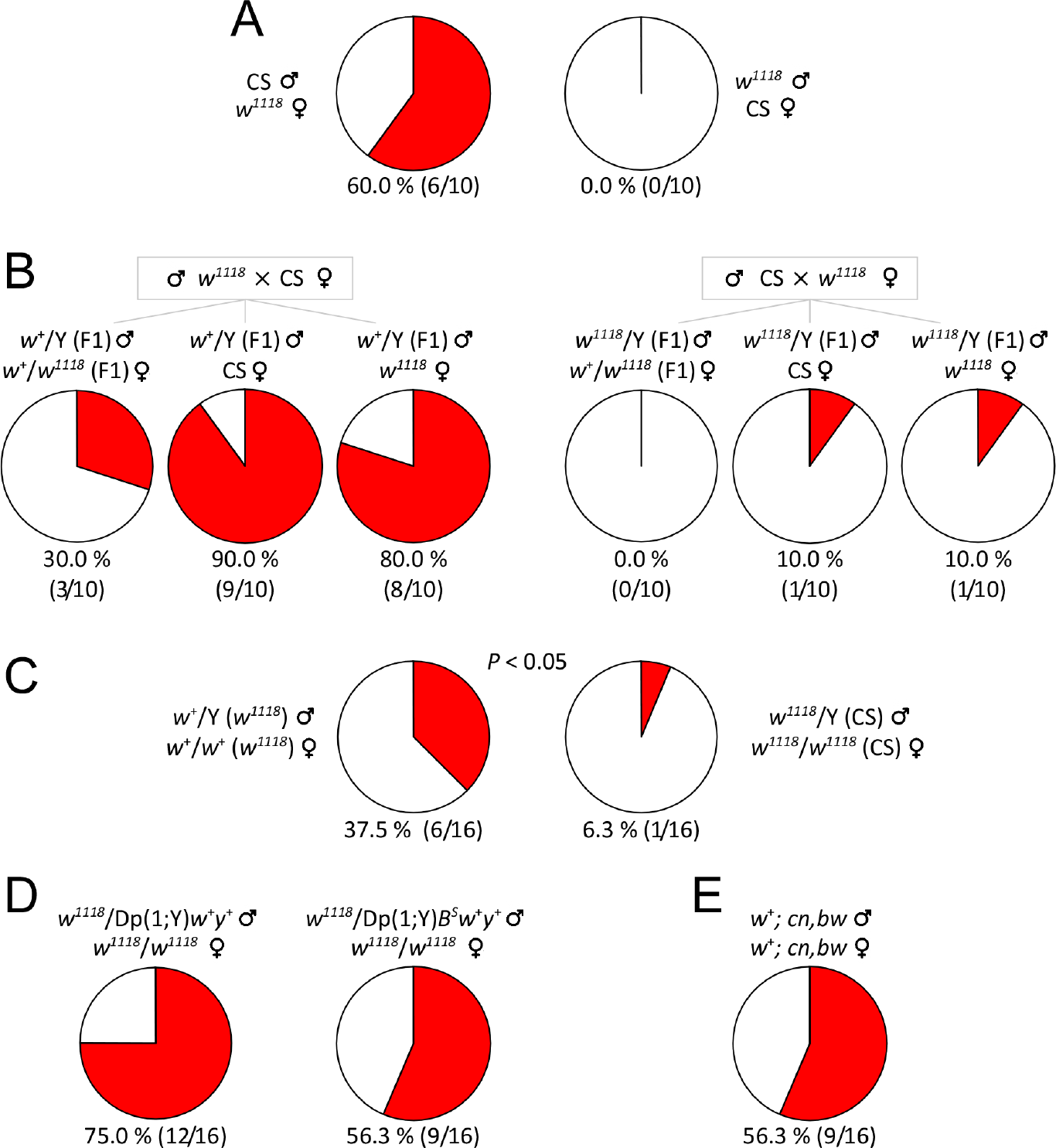
*w*^+^ allele was associated with copulation success. (**A**) Copulation success between CS males and *w*^1118^ females, and between *w*^1118^ males and CS females. (**B**) Copulation success between F1 males and three types of females (See Methods and text). (**C**) Copulation success of *w*^+^(*w*^1118^) and *w*^1118^(CS) flies in which different *w* alleles are exchanged between CS and *w*^1118^ by serial backcrossing for ten generations. *P* value from Fisher’s exact test. (**D**) Copulation success of flies with *w*^+^ duplicated to the Y chromosome. (**E**) Copulation success in *w*^+^; *cn,bw* (white-eyed) flies. For all the experiments, copulation success is indicated as red in the pie chart. Sexual activities of flies was observed within 60 minutes in the circular arenas. Numbers of tested pairs and numbers of successful copulation are shown. Fly sex is indicated as ♂ (male) and ♀ (female).

### *w*^+^ was associated with copulation success

*w*^1118^ fly carries a null allele of *w* on the X chromosome. We examined whether *w*^+^ was associated with copulation success. We first tested two different male progenies: *w*^+^/Y (F1) and *w*^1118^/Y (F1), produced by the cross between *w*^1118^ males and CS females, and the reciprocal cross between CS males and *w*^1118^ females. Male flies of each genotype were paired with three types of virgin females for the tests.

Paired with respective *w*^+^/*w*^1118^ sibling females, *w*^+^/Y (F1) showed a copulation success of 30.0% (3/10), whereas *w*^1118^/Y (F1) had no success (0.0%, 0/10) in copulation. Paired with CS females, *w*^+^/Y (F1) displayed significantly higher copulation success (90.0%, 9/10) than *w*^1118^/Y (F1) (10.0%, 1/10) (*P <* 0.05, Fisher’s exact test). Paired with *w*^1118^ females, *w*^+^/Y (F1) had higher copulation success (80.0%, 8/10) than *w*^1118^/Y (F1) (10.0%, 1/10) (*P <* 0.05, Fisher’s exact test) (Fig. 3B). Therefore, *w*^+^-carrying F1 males showed better copulation success than *w*^1118^-carrying F1 males.

We then examined copulation success of *w*^+^/Y (*w*^1118^) and *w*^1118^/Y (CS) flies in which different *w* alleles were exchanged between CS and *w*^1118^ flies by serial backcrossing for ten generations^34^. Copulation success between *w*^+^/Y (*w*^1118^) and sibling *w*^+^/*w*^+^ (*w*^1118^) females (37.5%, 6/16) was higher than that between *w*^1118^/Y (CS) and sibling *w*^1118^/*w*^1118^ (CS) females (6.3%, 1/16) (*P* = 0.0415, one-tailed Fisher’s exact test) (Fig. 3C). Results confirmed that *w*^+^-carrying males had increased copulation success compared with *w*^1118^-carrying males.

### *w*^+^ duplicated to Y chromosome rescued the defect of copulation success in *w*^1118^ flies

To further support the association between *w*^+^ and copulation success, we examined the copulation percentage of males carrying a *w*^+^ allele duplicated to the Y chromosome. Copulation success between *w*^1118^/Dp(1;Y)*w*^+^*y*^+^ males and their sibling females *w*^1118^/*w*^1118^ was 75.0% (12/16), which was comparable to CS (see Fig. 2E) with no significant difference (*P* = 0.71, Fisher’s exact test) (Fig. 3D). Similar data was obtained using another duplication line. Copulation success between *w*^1118^/Dp(1;Y)*B^S^w*^+^*y*^+^ males and sibling females *w*^1118^/*w*^1118^ (56.3%, 9/16) was comparable to CS with no significant difference (*P* = 0.08, Fisher’s exact test) (Fig. 3D). Thus, *w*^+^ duplicated to the Y chromosome rescued the defect of copulation success in *w*^1118^ flies.

### Phenotypic independence between copulation success and eye color

The classical phenotype for *w*^+^ is for red eye color. A critical question is whether the defect of copulation success might be due to the loss of normal eye color and the associated poor vision^35^. To test this possibility, we examined copulation success in *w*^+^; *cn*, *bw*, a white-eyed strain carrying *w*^+^ allele^24^. Copulation success was 56.3% (9/16) between *w*^+^; *cn*, *bw* males and sibling females (Fig. 3E). There was no statistical difference of copulation success between *w*^+^; *cn*, *bw* and CS (*P* = 0.08, Fisher’s exact test). Therefore, *w*^+^-associated copulation success was independent of eye color phenotype.

### m*w*^+^ rescued the defect of copulation success in *w*^1118^ flies

*Drosophila* m*w*^+^ is a miniature form of *w*^+^. m*w*^+^ has been widely used as a marker gene to indicate the genomic recombination of a transgene. Since *w*^+^ and m*w*^+^ encode the same protein, it becomes important to understand whether m*w*^+^ rescues the defect of copulation success in *w*^1118^ flies. We tested UAS lines with m*w*^+^-carrying transposons inserted into the X or autosomal (II or III) chromosome. All tested UAS flies were synchronized into the *w*^1118^ isogenic background. The UAS but not Ga^l4^ or other transgenic flies were chosen in order to minimize possible complex effects due to ectopic expression of a transcription factor in addition to m*w*^+24^.

Paired with CS virgin females, male flies with one copy of m*w*^+^ on the autosome showed 0.0 - 12.5% copulation success (Fig. 4A). Males with m*w*^+^ on the X chromosome displayed 6.3 - 62.5% copulation success (Fig. 4B). Therefore, males with one copy of m*w*^+^ showed an increase of copulation success relative to *w*^1118^ males. The observed variance among different UAS lines could be due to different expression of m*w*^+^ or potentially a second mutation. Likely, the rescue effect of m*w*^+^ was strong if integrated on the X chromosome, perhaps because of dosage compensation for genes on the X chromosome^36, 37^.

**Figure 4.**
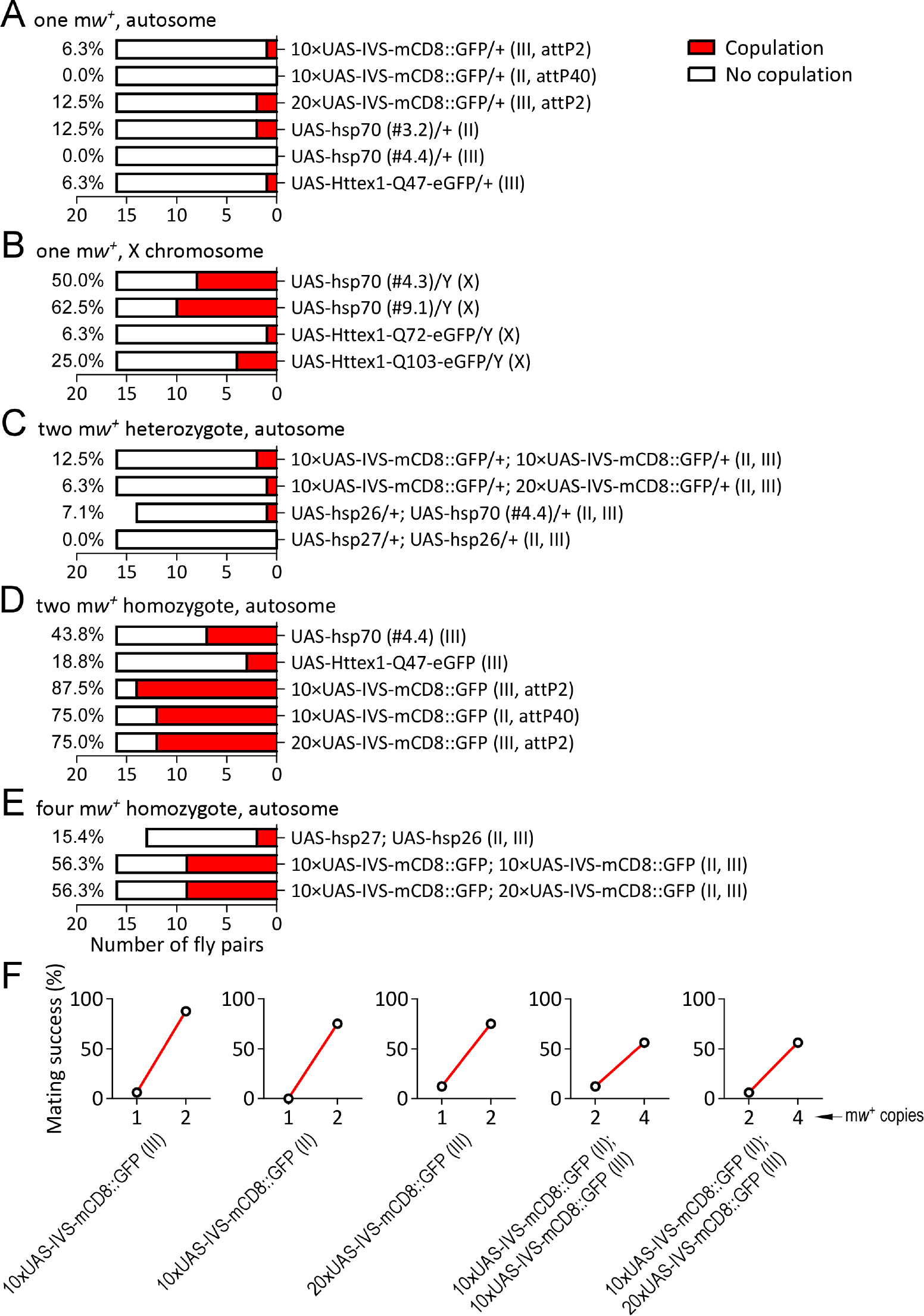
m*w*^+^ rescued the defect of copulation success in *w*^1118^ flies. (**A**) Copulation success (red bar) in flies carrying one copy of m*w*^+^ on the autosome. The X axis denotes the number of fly pairs (sample size). Values of percent copulation success are provided. (**B**) Copulation success in flies carrying one copy of m*w*^+^ on the X chromosome. (**C**) Copulation success in flies carrying two heterozygous m*w*^+^ (two m*w*^+^ heterozygotes) on the autosomes. (**D**) Copulation success in flies carrying homozygous m*w*^+^ (two m*w*^+^ homozygotes) on the autosomes. (**E**) Copulation success in flies carrying double homozygous m*w*^+^ (four m*w*^+^ homozygotes) on the autosomes. (**F**) Correlations of copulation success between heterozygous and homozygous flies carrying the same transposons. Numbers indicate m*w*^+^ copies on the autosomes. Chromosomal locations of m*w*^+^ are indicated in the parentheses. attP2: site-specific recombination site on the third chromosome; attP40: site-specific recombination site on the second chromosome.

Males carrying two copies of m*w*^+^ (heterozygous for each) on the autosomes displayed 0.0 - 12.5% copulation success (Fig. 4C). Males with homozygous m*w*^+^ (two copies) on the autosomes displayed 18.5 - 87.5% copulation success (Fig. 4D). Males with two homozygous m*w*^+^ (four copies) on the autosomes showed 15.4 - 56.3% copulation success (Fig. 4E). These data supported the conclusion that m*w*^+^ rescued the defect of copulation success in *w*^1118^ males.

Notably, males carrying homozygous m*w*^+^ alleles had increased copulation success compared with males carrying heterozygous m*w*^+^. Copulation success was increased from 6.3% for heterozygous to 87.5*%* for homozygous 10×UAS-IVS-mCD8::GFP (on III) flies (Fig. 4F). Similar results were observed as 0.0*%* to 75.0*%* in 10×UAS-IVS-mCD8::GFP (on II), 12.5*%* to 75.0*%* in 20×UAS-IVS-mCD8::GFP (on III), 12.5*%* to 56.3*%* in 10×UAS-IVS-mCD8::GFP;10×UAS-IVS-mCD8::GFP (on II and III), 6.3*%* to 56.3*%* in 10×UAS-IVS-mCD8::GFP;20×UAS-IVS-mCD8::GFP (on II and III), 0.0*%* to 43.8*%* in UAS-hsp70#4.4 (on III), 0.0*%* to 15.4*%* in UAS-hsp27;UAS-hsp26 (on II and III), and 6.3*%* to 18.8*%* in UAS-Httex1-Q47-eGFP (on III) (Fig. 4A and 4C-F). The increase of copulation success for each UAS line was observed from flies with heterozygous m*w*^+^ allele to flies with homozygous alleles carried in the same transposon. Hence they would have different levels of White protein with the same expression pattern under the same genetic background. There was a strong positive correlation between copies of m*w*^+^ and the percentage of copulation success. These data indicate that m*w*^+^ rescued the defect of copulation success in a dosage-dependent manner.

### m*w*^+^ rescued courtship rituals in *w*^1118^ flies

A wild-type male often attempts a series of courtship ritual towards a female before copulation^2, 3^. We examined whether m*w*^+^ rescued the reduced courtship in white-eyed flies^28^. A courtship index was calculated as the fraction of time during which observable courtship activities (including orientation, female following and wing extension) occur^38^.

Courtship indices were calculated in CS, *w*^1118^ and m*w*^+^-carrying UAS flies. During the first three minutes, most CS males displayed typical courtship behaviors to CS females, whereas w1118 males showed greatly reduced courting activities. Heterozygous 10×UAS-IVS-mCD8::GFP (III) flies (m*w*^+^/+) displayed sporadic courting, while homozygous flies (m*w*^+^/m*w*^+^) showed strong and persistent courting (Fig. 5A).

**Figure 5.**
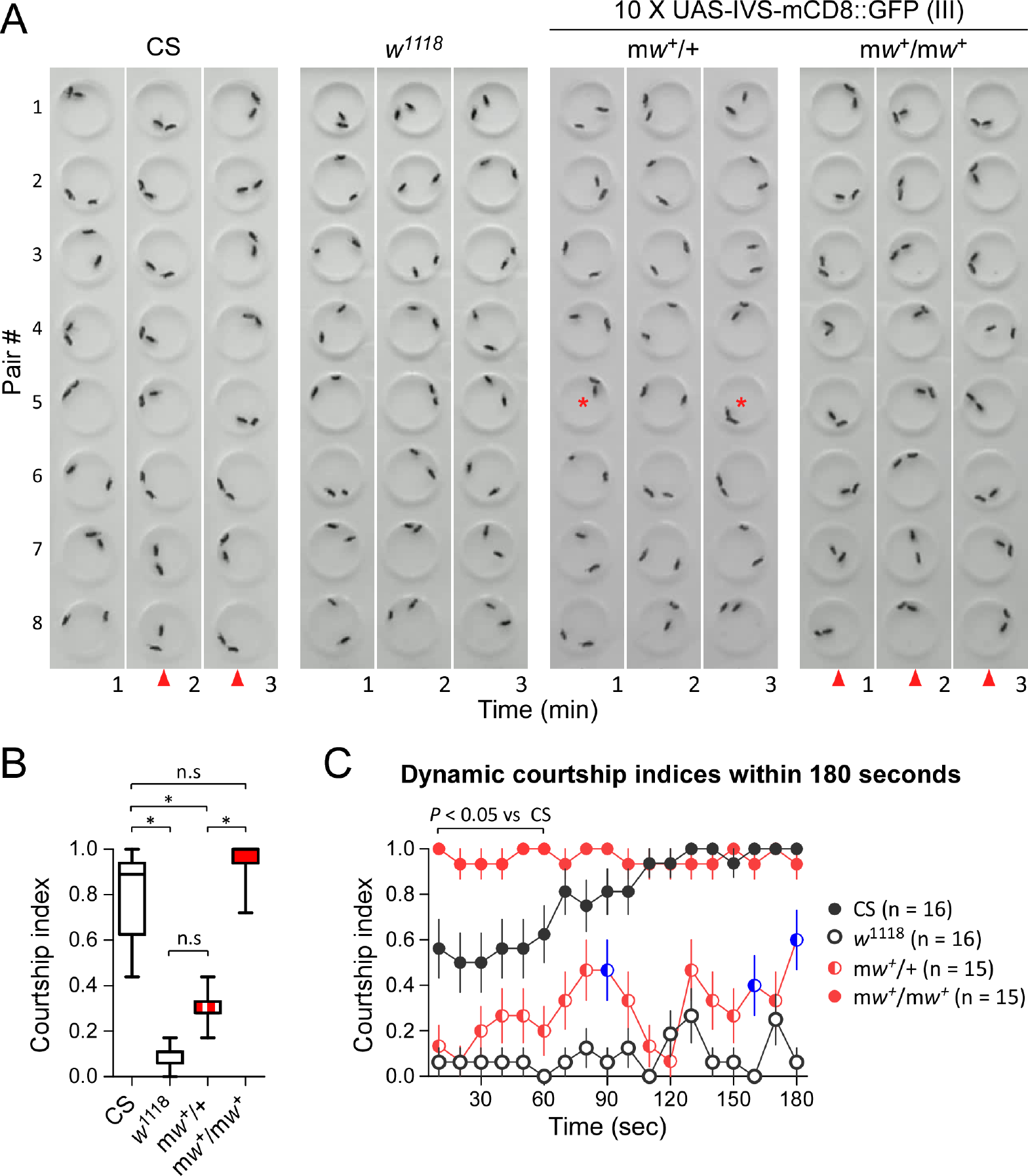
m*w*^+^ rescued the reduced courtship in *w*^1118^ flies. (A) Courtship activities before copulation in CS, *w*^1118^ and m*w*^+^ flies. Shown are the video-frames sampled once per minute during the first three minutes. Consistent courting between pairs at a time point in all or most pairs (red triangle), and sporadic courting seen in individual arenas (red star) are indicated. There are eight pairs for each genotype. Each pair contains a naive male and a virgin female. (**B**) Courtship indices within three minutes in tested flies. *, *P <* 0.05 (one-way ANOVA); n.s., non-significance. (**C**) Dynamic courtship indices within the first 180 seconds in four genotypes. Courtship indices were repeatedly evaluated once every 10 seconds. *P* value is from two-way ANOVA. Sample sizes are indicated.

Within three minutes, the median courtship index of CS males was 0.83 (IQR 0.63–0.94, n = 16), *w*^1118^ 0.11 (IQR 0.06–0.11, n = 16), heterozygous m*w*^+^ 0.28 (IQR 0.28–0.33, n = 15) and homozygous m*w*^+^ 1.00 (IQR 0.94–1.00, n = 15). Male *w*^1118^ showed a markedly reduced courtship index compared with CS (*P <* 0.05, Kruskal-Wallis test with Dunn’s multiple comparisons). Heterozygous m*w*^+^ males displayed a slightly increased courtship index compared with *w*^1118^ males with insignificant difference, but a reduced courtship index compared with CS (*P <* 0.05, Kruskal-Wallis test with Dunn’s multiple comparisons). Homozygous m*w*^+^ males showed a courtship index higher than heterozygous m*w*^+^ (*P <* 0.05, Kruskal-Wallis test with Dunn’s multiple comparisons) (Fig. 5B). Interestingly, Homozygous m*w*^+^ males had a courtship index similar to CS males (with not significant difference). Therefore, m*w*^+^ rescued the reduced courtship activity in *w*^1118^ males, and the rescue effect was mw+ copy number-dependent.

We further examined the dynamic changes of courtship indices in CS, *w*^1118^ and m*w*^+^ flies within the first 180 seconds. The initial courtship index (during the first ten seconds) was around 0.6 for CS males and around 0.1 for *w*^1118^ males, indicating rapid engagement of courtship with females in most CS flies but extremely slow in *w*^1118^ flies. The initial courtship index was around 0.1 for heterozygous m*w*^+^ males and 1.0 for homozygous m*w*^+^ males, thus almost all homozygous m*w*^+^ males but not heterozygous males started courtship immediately. CS males had an average courtship index of 0.5–0.6 in the first 60 seconds and a gradual increase up to nearly 1.0 within the next 120 seconds, whereas *w*^1118^ males had a courtship index of 0.0–0.2 most of the time. Courtship indices of CS males were higher than those of *w*^1118^ males throughout the observation duration (*P <* 0.01, repeated measures ANOVA with Bonferroni post-tests). Heterozygous m*w*^+^ flies showed increased courtship indices in some of the observation periods (i.e. 90, 160 and 180 sec) compared with *w*^1118^ flies, but reduced levels in most of the observations compared with CS (repeated measures ANOVA with Bonferroni post-tests). Homozygous m*w*^+^ males had a persistently high courtship index (0.9–1.0) throughout, and surprisingly, during the first 60 seconds, courtship indices of homozygous m*w*^+^ males were even higher than CS males (*P <* 0.05, repeated measures ANOVA with Bonferroni post-tests) (Fig. 5C). These data suggest that m*w*^+^ rescued the reduced courtship activity in *w*^1118^ flies, and that homozygous m*w*^+^ alleles over-rectified the courtship in *w*^1118^ males.

### Male-female sexual experience rectified the defect of copulation success in *w*^1118^ flies

To understand how *w*^1118^ males might have lost copulation ability in the circular arenas, we attempted to promote mating of *w*^1118^ flies through several approaches: (1) increase of arena size, (2) dim red illumination and (3) sexual training.

In small arenas flies often become more active in behavior and locomotion than large arenas^3, 34^. The copulation success of *w*^1118^ could be suppressed by spatial restriction. CS and *w*^1118^ males reduce locomotion in large arenas (3.81 cm diameter) compared with small ones (1.27 cm diameter)34, suggesting reduced disturbance of behavior in large arenas. Copulation success of CS pairs in large arenas was 66.7% (6/9), a level similar to that in small arenas (see Fig. 2E) (*P >* 0.05, Fisher’s exact test). There was no copulation observed in *w*^1118^ pairs (0.0%, 0/9) (Fig. 6A). Thus, the increase of arena size had no effect on copulation success in wild-type, and failed to promote copulation in *w*^1118^ flies.

**Figure 6.**
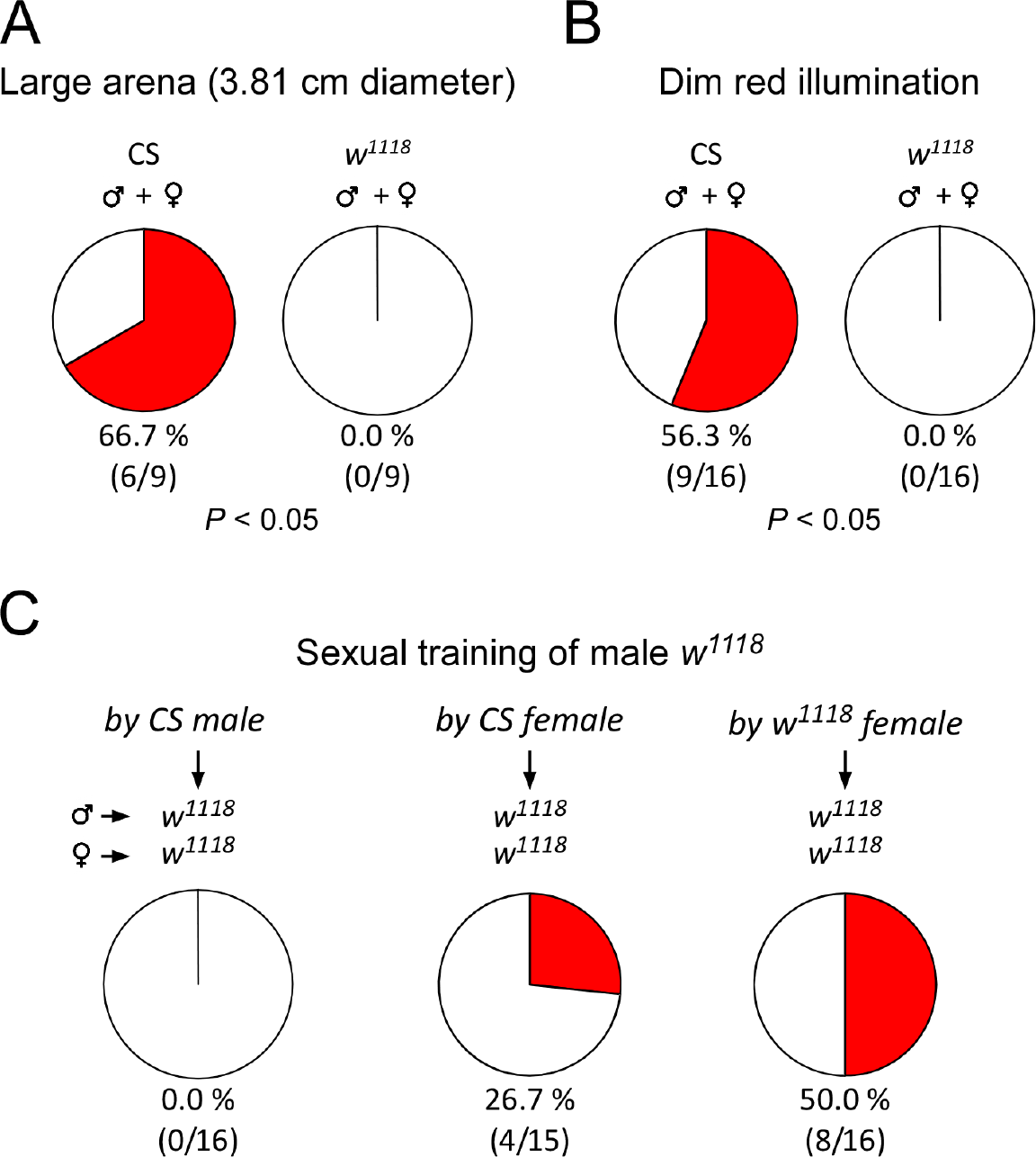
Male-female sexual experience promoted copulation success in *w*^1118^ flies. (**A**) Copulation success (red) of CS and *w*^1118^ flies in large arenas (3.81 cm diameter 0.3 cm depth). *P* value is from Fisher’s exact test. (**B**) Copulation success of CS and *w*^1118^ flies under dim red illumination. *P* value is from Fisher’s exact test. (**C**) Copulation success between sexually trained *w*^1118^ males and *w*^1118^ virgin females. Training conditions are indicated (see Methods for description).

Daylight illumination reduces sexual arousal in *w*^1118^ males^28^. Diminished light improves copulation in *ebony* and *lozenge*^3^ flies^10, 39^. It is possible that copulation success of *w*^1118^ increases under dim red illumination, a condition mimicking darkness due to poor visual sensitivity to red light^40^. With dim red illumination, copulation success was 56.3% (9/16) in CS pairs and 0.0% (0/16) in *w*^1118^ flies (Fig. 6B). There was clearly a defect of copulation success in *w*^1118^ compared with CS (*P <* 0.05, Fisher’s exact test). Dim red illumination did not repair the defect of copulation success in *w*^1118^ flies.

Mating behavior can be remodeled by male-male courtship^41–43^. Specific ratios of white-eyed : wild-type males (i.e. *<* 0.4, or *>* 0.8) improve copulation success in white-eyed flies, if two strains (including white-eyed males and females, and wild-type males and females) are present together^33^. We examined whether *w*^1118^ males could learn to improve copulation success from CS males. Naive *w*^1118^ males and CS males were mixed at 9:1 and raised for four days. Paired with *w*^1118^ virgin females, *w*^1118^ males showed no copulation (0.0%, 0/16) in the circular arenas (Fig. 6C).

Male-female copulation experience enhances mating behavior and success in wild-type flies^44^. Male-female sexual training might be more effective than male-male courtship training in improving copulation success in *w*^1118^ flies. Naive *w*^1118^ males and CS virgin females were mixed with equal numbers and raised for four days. Paired with four-day-old *w*^1118^ virgin females, experienced *w*^1118^ males showed 26.7% (4/15) copulation success. If naive *w*^1118^ males were sexually experienced with *w*^1118^ virgin females, copulation success in *w*^1118^ flies was increased to 50.0% (8/16) (Fig. 6C), which was significantly higher than that in non-experienced *w*^1118^ flies (see Fig. 2E) (*P <* 0.05, Fisher’s exact test). The male-female sexual training for four days always resulted in the production of offspring, indicating the occurrence of copulation during training. Therefore, male-female copulation experience rectified the defect of copulation success during arena trials in *w*^1118^ flies.

## Discussion

*Drosophila* male-female copulation following the courtship ensures a physical contact of reproductive systems for the transfer of genetic contents to the offspring. There is a clear involvement of *w* in sexual discrimination^2, 26^ and courtship^12, 13^ in male flies, but whether *w*^+^ determines male-female copulation success is unknown. Here we show that loss-of-*w* is associated with a defect of copulation success in a circular arena, that *w*^+^-associated copulation success is independent of eye color phenotype, and that addition of m*w*^+^ into a null background rescues male-female copulation success in a manner that is m*w*^+^ copy number-dependent. These data add consolidated genetic evidence that *w*^+^ controls male-female copulation success in *Drosophila melanogaster*.

Rapid engagement of courtship is evident between a wild-type naive male and a wild-type virgin female in a circular arena. In contrast, extremely low chance of courtship is observed between a white-eyed naive male and a white-eyed virgin female. The small size of arena (1.27 cm diameter) induces increased locomotion in males of both wild-type and white-eyed mutant^34^. However, once a virgin female is present, a wild-type male rapidly switches the behavior from locomotion to courting, as revealed by a high initial courtship index. These observations suggest that there is a high priority for courtship relative to exploratory activity in the small arenas in wild-type flies. Indeed, copulation behavior in wild-type is vigorous and highly resistant to environmental stress such as a strong smell of ether^2^. We also find that the change of arena size does not affect copulation success in wild-type. On the other hand, naive white-eyed males do not show rapid engagement of courting, and fail to copulate with virgin females in the arenas. Thus, a high priority for copulation might have been lost or severely impaired in white-eyed flies.

We offer here two main reasons that could explain the defect of copulation success of white-eyed flies in the circular arenas. First, *w*^+^ directly controls male-female copulation success. Second, loss-of-*w*^+^ delays sexual development and maturation, resulting in impaired male-female copulation success.

Addition of *w*^+^ to white-eyed flies by exchanging the allele to the X chromosome or duplicating to the Y chromosome rescues the defect of copulation success. These findings indicate a strong association between *w*^+^ and copulation success. This is further supported by the observations that addition of m*w*^+^ to the *w* null background rectifies the defect of copulation success, and that the rectification is m*w*^+^ copy number-dependent. An added value of m*w*^+^ copy number-dependent rectification is the practical approach for genetic manipulation of copulation success by m*w*^+^ allele. By inserting m*w*^+^ to the genome and varying the copy number, we are able to control male-female copulation success in a manner that is dosage-dependent. These findings suggest that *w*^+^ directly controls male-female copulation success.

A four-day-old wild-type naive male could have highly accumulated sex drive^2^, whereas a four-day-old wild-type virgin female might have reduced receptiveness^45^. Brought together, they show a level of copulation success comparable to previous observations in several wild-types^29–32^. Thus, sexual isolation for four days would have no or little impact to male-female copulation success in wild-type. However, sexual isolation of white-eyed flies, particularly males, does have a significant impact to copulation success. This is evident that sexual training of white-eyed males by sibling virgin females promotes copulation success between trained males with isolated virgin females. Because male-male training from wild-types is unsuccessful, the training has to be with females and apparently, involves male-female copulation experience, as judged by the production of offspring. These data are consistent with the observation that male-female copulation confers competitive advantages for copulation success in wild-type males^44^. The specific requirement of sexual training is different from that used for *fru^M^* mutant, which improves courtship behavior by the training of any sex of flies^46^. Thus, male-female copulation experience is critical for white-eyed males to achieve copulation success. Such a male-female copulation experience is, however, not required for wild-type flies and most tested m*w*^+^ flies for successful copulation. Taking into account that *w*^+^ has an implicated role for learning and memory23, naive white-eyed flies might have impaired ability to learn or develop normal sexual performance, and this impairment could be rectified by sexual training or learning. We therefore explain our findings as that loss-of-*w*^+^ delays sexual development and maturation, resulting in impaired male-female copulation success.

Much effort has been made to understand *Drosophila* courtship behavior in a single sex (e.g. male fly). Little is known about the genetic basis for *Drosophila* male-female copulation success. Our findings extend currently existing understanding of *w*^+^/m*w*^+^ on courtship behavior in several aspects.

It is novel that *w*^+^-associated copulation success is independent of eye color phenotype. White-eyed flies have poor visual acuity, which affects optomotor response^35^, exploratory behavior^47^, and mating competition under light conditions^48^. Those white-eyed flies carry *w*, a null mutant allele. It is reasonable to believe that both phenotypes, copulation success and eye color, are tightly linked to each other and could have been lost at the same time due to the loss-of-*w*^+^. Here we show that these two phenotypes are clearly separable. By blocking pteridine and ommochrome pigment pathways^49, 50^, we generate a white-eyed strain carrying *w*^+^. These *w*^+^-carrying white-eyed flies show wild-type-like copulation success in the circular arenas. Our findings clarify two concerns. The white-eye phenotype and its associated visual acuity do not necessarily affect male-female copulation success. And, *w*^+^ possesses clearly separable functions. It has been shown that *w*^+^ promotes fast locomotor recovery from anoxia, another phenotype independent of eye color^24, 25, 51^. Therefore, we provide solid evidence to support a pleiotropic function of *w*^+^ in controlling male-female copulation success.

Second, we highlight that m*w*^+^ causes abnormal male-female copulation success in addition to abnormal courtship. Wild-type male and female flies have tightly regulated expression of *w*^+^, and do not require *w*^+^-copy-number-dependent copulation success. That m*w*^+^-copy-number-dependent copulation success is apparently an abnormal phenomenon in *Drosophila*. Also, the incomplete rescue of courtship index by heterozygous m*w*^+^ and over-rectification of courtship index by homozygous m*w*^+^ indicate the abnormal recovery of courtship behavior. The commonly observed variation of eye color in m*w*^+^-carrying flies is another example of abnormal recovery of *w*^+^ function.

Third, we examine a clearly observable phenotype, male-female copulation success, which involves both fly sexes. Behavioral characteristics of male-female copulation were described in 1910s^2^. Using advanced genetic resources (e.g. m*w*^+^-carrying flies), we provide genetic evidence that *w*^+^ controls male-female copulation success. Additionally, this phenotype differs from courtship, a sexual request occurring at stages of pre- and post-copulation. Successful copulation, if observed in our laboratory settings, occurs only once within a period of 3-hr. However, the courtship of a male happens at multiple stages, even after the completion of copulation towards the mated female (personal observations). Thus, male-female copulation success cannot be evaluated by courtship activity.

How might *w*^+^ control male-female copulation success? Since the first report more than 100 years ago^16^, it has long been believed that *w*^+^ product transports and deposits pigment in pigmental cells in *Drosophila*^52, 53^. It was not until 1995 that an extra-retinal neural function of *w*^+^ was proposed^12^. Since then studies have shown that the White protein uptakes biogenic amines, second messengers, intermediate metabolites and many small molecules including pigment precursors into vesicles/granules and transports to appropriate subcellular locations^15, 19, 20, 22, 52^. The behavioral performance of wild-type flies supports the proposed neural function of *w*^+^. Wild-type flies have a fast reaction to volatile general anesthetics (VGAs)54, fast and consistent locomotor recovery from anoxic coma^24, 55^, and high boundary preference of exploratory activities in the circular arenas^34^. In contrast, loss-of-*w* mutants show altered and light-dependent sensitivity to VGAs^54^, delayed locomotor recovery from anoxia^24^, reduced boundary preference of exploration^34^, and reduced learning and memory of thermal stress^23^. Therefore, it is highly plausible that the White protein possesses housekeeping functions in maintaining appropriate vesicular contents, transporting molecular substrates, and improving signaling efficacy in the nervous system. These housekeeping functions of White could be essential for male-female copulation success.

## Methods

### Flies

Fly strains used in the current study and their sources are listed in Table 1. Flies were maintained with standard medium (cornmeal, agar, molasses and yeast) at 21-23 °C in a light/dark (12/12 hr) condition. *w*^1118^(CS) and *w*^+^(*w*^1118^) flies carry different *w* alleles exchanged between CS and *w*^1118^ flies by serial backcrossing for ten generations^34^. The *w*^+^ duplication (to the Y chromosome) line *w*^1118^/Dp(1;Y)*B^S^w*^+^*y*^+^ was derived from Raf^11^/FM6, l(1)FMa^1^/Dp(1;Y)*B^S^w*^+^*y*^+^ (Bloomington *Drosophila* Stock Center (BDSC) #5733) by crossing a single male with three *w*^1118^ females, and their male progeny into *w*^1118^ females to establish a stock^24^. In this study we generated another *w*^+^ duplication line *w*^1118^/Dp(1;Y)*w*^+^*y*^+^ from dwg^11*−*32^/Dp(1;Y)*w*^+^*y*^+^/C(1)DX, *y*^1^*f*^1^ (BDSC #7060). UAS-hsp2656, UAS-hsp2756, UAS-hsp70 (#3.2, #4.3, #4.4 and #9.1)^57^ and UAS-Httex1-Qn-eGFP (n = 47, 72 or 103)^58^ were generated under *w*^1118^ genetic background. 10×UAS-IVS-mCD8::GFP (at attP2 or attP40, BDSC #32185, #32186) and 20×UAS-IVS-mCD8::GFP (attP2, BDSC #32194) flies were backcrossed to *w*^1118^ background for ten generations. Flies with combinations of UAS-IVS-mCD8::GFP were generated after the backcrossing.

**Table 1.**
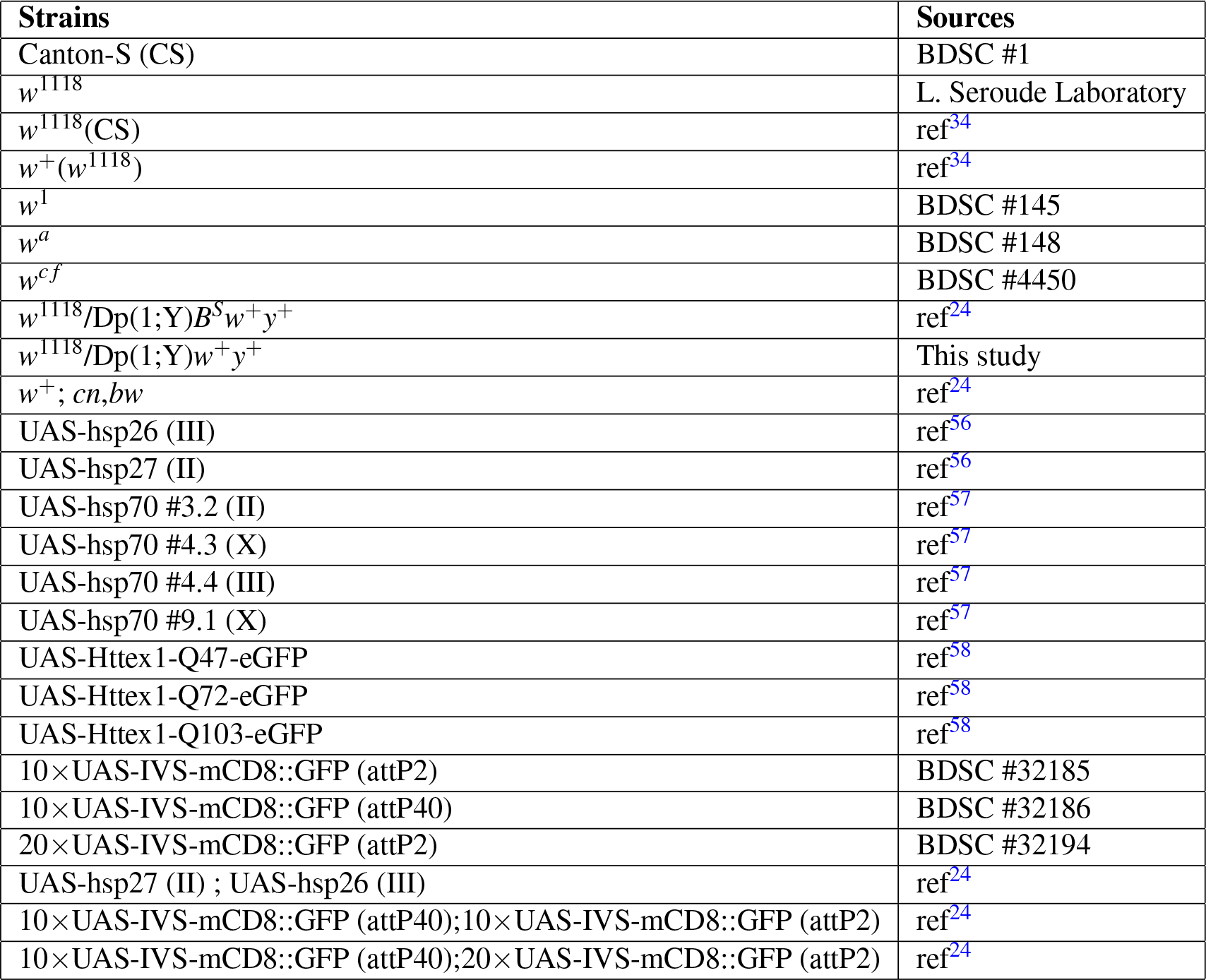
Fly strains used in this study and their sources. BDSC, Bloomington *Drosophila* Stock Center; attP2, Site-specific recombination site in the third chromosome; attP40, Site-specific recombination site in the second chromosome; ref, reference.

### Fly preparation for copulation analysis

Naive males and virgin females were collected within five hours after eclosion. Single-sexed flies were grouped as 10-20 per vial, and aged to 4-7 days before the experiments. We used nitrogen gas to knock down flies during collection time. Tested flies were free of nitrogen exposure for at least three days since the initial collection.

Sexual training of *w*^1118^ males was carried out as follows: (1) male-male courtship training by CS males: 18 naive *w*^1118^ males and 2 naive CS males were mixed in each vial and raised for four days. Male flies obtain sexual experience by courting naive males^41–43^. The ratio was determined based on a report that white-eyed males rare or predominant to wild-type males in a mixed population increase copulation success with white-eyed females^33^; (2) training by CS females: 10 naive *w*^1118^ males and 10 virgin CS females were mixed in each vial and raised for four days; (3) training by *w*^1118^ females: 10 naive *w*^1118^ males and 10 virgin *w*^1118^ females were mixed in each vial and raised for four days. Trained *w*^1118^ males were then paired individually with a four-day-old *w*^1118^ virgin female for copulation analysis. Male-female copulation has occurred in every type of male-female training because of the observed production of offspring.

### Analysis of copulation success

The apparatus for copulation observation was the same as previously reported^34^. Briefly, a naive*/*trained male and a virgin female were gently aspirated into a circular arena (1.27 cm diameter 0.3 cm depth). Sexual behavior was monitored and video-captured with a digital camera (Webcam C905, Logitech) for 60 minutes or a duration indicated otherwise. Copulation success (defined as apparent physical attachment between a male and a female) was post-analyzed from videos. The percentage of copulation success for each genotype was examined from a minimum of nine pairs. In many experiments the sibling male and female were used for the analysis in order to avoid sexual reluctance between heterospecific flies^59, 60^ or potential reluctance between different strains. In several experiments uniform females (e.g. CS females) or females of different strains were used. Specific genotypes of flies are described in the text. Experiments were conducted during the light time and at least three hours away from light-dark transit. Illumination was provided with a light box (Logan portaview slide/transparency viewer) with surrounding reflections. Dim red illumination was generated by using a red filter (600 nm long-pass, Roscolux #26, Rosco Canada) on the light box.

### Evaluation of male courtship activity

The courtship behavior of a male fly before copulation was evaluated using a courtship index38. In general, sexual activities of fly pairs were sampled once every 10 seconds during the first 180 seconds. At every sampling time, if for each pair the courtship activity was observed a score was assigned as 1, otherwise 0. A courtship index was calculated as the fraction of observation time during which any courtship happened. Clearly observable courtship activities of males included orientation, female following, and wing extension. We chose the first 180 seconds for the evaluation of courtship because copulation did not occur within this period in most of the tested flies.

### Statistics

Fisher’s exact test was used to examine the percentage of copulation success between two different strains. Kruskal-Wallis test with Dunn’s multiple comparisons was performed to analyze courtship index between different groups of flies. Two-way ANOVA with Bonferroni post-tests was conducted to examine dynamic courtship index during the first 180 seconds. Data with normal distribution were presented as average ± standard error of mean (mean ± SEM). Data with non-normal distribution were illustrated as box-plots. A *P <* 0.05 was considered significant difference.

## Acknowledgements

This work was funded by Natural Sciences and Engineering Research Council of Canada (NSERC) grant (RGPIN 40930-09) to R.M.R.

## Author contributions statement

C.X. conceived the experiments; C.X. and S.Q. conducted the experiments, analysed the data, and prepared the manuscript; R.M.R. provided funding support and manuscript editing. All authors reviewed the manuscript.

## Additional information

The authors declare no conflict of interest.

## References

1. Mayr, E. Experiments on sexual isolation in *Drosophila* VII. The nature of the isolating mechanisms between *Drosophila pseudoobscura* and *Drosophila persimilis*. Proc Natl Acad Sci U S A 32, 128–137 (1946).

2. Sturtevant, A. H. Experiments on sex recognition and the problem of sexual selection in *Drosoophilia*. J. Animal Behav. 5, 351–366 (1915). DOI 10.1037/h0074109.

3. Spieth, H. T. Mating behavior within the genus *Drosophila* (diptera). Bull Am Mus Nat Hist 99, 395–474 (1952).

4. Miller, D. D. Sexual isolation and variation in mating behavior within *Drosophila athabasca*. Evol. 12, 72–81 (1958).

5. Spiess, E. B., Langer, B. & Li, C. Chromosomal adaptive polymorphism in *Drosophila persimilis* III. Mating propensity of homokaryotypes. Evol. 15, 535–44 (1961).

6. Spiess, E. B. & Langer, B. Mating speed control by gene arrangements in *Drosophila pseudoobscura* homokaryotypes. Proc Natl Acad Sci U S A 51, 1015–1019 (1964).

7. Miller, D. D. & Westphal, N. J. Further evidence on sexual isolation within *Drosophila athabasca*. Evol. 21, 479–92 (1967).

8. Patty, R. A. Investigation of genetic factors influencing duration of copulation in ‘eastern’and ‘western’ Drosophila athabasca. Anim Behav 23, 344–348 (1975).

9. Merrell, D. J. Selective mating in *Drosophila melanogaster*. Genet. 34, 370–389 (1949).

10. Jacobs, M. E. Influence of light on mating of *Drosophila melanogaster*. Ecol. 41, 182–188 (1960). DOI 10.2307/1931952.

11. Connolly, K., Burnet, B. & Sewell, D. Selective mating and eye pigmentation: an analysis of the visual component in the courtship behavior of *Drosophila melanogaster*. Evol. 23, 548–59 (1969).

12. Zhang, S. D. & Odenwald, W. F. Misexpression of the *white (w)* gene triggers male-male courtship in *Drosophila*. Proc Natl Acad Sci U S A 92, 5525–9 (1995).

13. Hing, A. L. & Carlson, J. R. Male-male courtship behavior induced by ectopic expression of the *Drosophila white* gene: role of sensory function and age. J Neurobiol 30, 454–64 (1996). DOI 10.1002/(SICI)1097-4695(199608)30:4¡454::AID-NEU2¿3.0.CO;2-2.

14. Hirai, Y., Sasaki, H. & Kimura, M. T. Copulation duration and its genetic control in *Drosophila elegans*. Zool Sci 16, 211–214 (1999).

15. Anaka, M. et al. The *white* gene of *Drosophila melanogaster* encodes a protein with a role in courtship behavior. J Neurogenet 22, 243–76 (2008). DOI 10.1080/01677060802309629.

16. Morgan, T. H. Sex limited inheritance in *Drosophila*. Sci. 32, 120–2 (1910). DOI 10.1126/science.32.812.120.

17. O’Hare, K., Murphy, C., Levis, R. & Rubin, G. M. DNA sequence of the *white* locus of *Drosophila melanogaster*. J Mol Biol 180, 437–55 (1984).

18. Hazelrigg, T. The *Drosophila white* gene: a molecular update. Trends Genet. 3, 43–7 (1987). DOI 10.1016/0168-9525(87)90165-x.

19. Sullivan, D. T. & Sullivan, M. C. Transport defects as the physiological basis for eye color mutants of *Drosophila melanogaster*. Biochem. Genet. 13, 603–13 (1975).

20. Sullivan, D. T., Bell, L. A., Paton, D. R. & Sullivan, M. C. Purine transport by malpighian tubules of pteridine-deficient eye color mutants of *Drosophila melanogaster*. Biochem. Genet. 17, 565–73 (1979).

21. Borycz, J., Borycz, J. A., Kubo´w, A., Lloyd, V. & Meinertzhagen, I. A. *Drosophila ABC* transporter mutants *white, brown* and *scarlet* have altered contents and distribution of biogenic amines in the brain. J Exp Biol 211, 3454–66 (2008). DOI 10.1242/jeb.021162.

22. Evans, J. M., Day, J. P., Cabrero, P., Dow, J. A. & Davies, S. A. A new role for a classical gene: *white* transports cyclic GMP. J Exp Biol 211, 890–9 (2008). DOI 10.1242/jeb.014837.

23. Sitaraman, D. et al. Serotonin is necessary for place memory in *Drosophila*. Proc Natl Acad Sci U S A 105, 5579–84 (2008). DOI 10.1073/pnas.0710168105.

24. Xiao, C. & Robertson, R. M. Timing of locomotor recovery from anoxia modulated by the *white* gene in *Drosophila*. Genet. 203, 787–797 (2016). DOI 10.1534/genetics.115.185066.

25. Xiao, C. & Robertson, R. M. White - cGMP interaction promotes fast locomotor recovery from anoxia in adult *Drosophila*. PLoS One 12, e0168361 (2017). DOI 10.1371/journal.pone.0168361.

26. Reed, S. & Reed, E. Natural selection in laboratory populations of *Drosophila*. II. Competition between a white-eye gene and its wild type allele. Evol. 4, 34–42 (1950).

27. Nilsson, E. E. et al. *Fruitless* is in the regulatory pathway by which ectopic *mini-white* and transformer induce bisexual courtship in *Drosophila*. J Neurogenet 13, 213–32 (2000).

28. Krstic, D., Boll, W. & Noll, M. Influence of the *white* locus on the courtship behavior of *Drosophila* males. PLoS One 8, e77904 (2013). DOI 10.1371/journal.pone.0077904.

29. MacBean, I. T. & Parsons, P. A. Directional selection for duration of copulation in *Drosophila melanogaster*. Genet. 56, 233 (1967).

30. Merrell, D. J. Mating between two strains of *Drosophila melanogaster*. Evol. 3, 266–268 (1949).

31. Spiess, E. B. Courtship and mating time in *Drosophila pseudoobscura*. Anim Behav 16, 470–9 (1968).

32. Ito, H. et al. Sexual orientation in *Drosophila* is altered by the satori mutation in the sex-determination gene fruitless that encodes a zinc finger protein with a BTB domain. Proc Natl Acad Sci USA 93, 9687–9692 (1996).

33. Ehrman, L. Mating success and genotype frequency in *Drosophila*. Anim Behav 14, 332–9 (1966).

34. Xiao, C. & Robertson, R. M. Locomotion induced by spatial restriction in adult *Drosophila*. PLoS One 10, e0135825 (2015). DOI 10.1371/journal.pone.0135825.

35. Kalmus, H. The optomotor responses of some eye mutants of *Drosophila*. J Genet. 45, 206–213 (1943).

36. Qian, S. & Pirrotta, V. Dosage compensation of the *Drosophila white* gene requires both the X chromosome environment and multiple intragenic elements. Genet. 139, 733–44 (1995).

37. Arkhipova, I., Li, J. & Meselson, M. On the mode of gene-dosage compensation in *Drosophila*. Genet. 145, 729–736 (1997).

38. Siegel, R. W. & Hall, J. C. Conditioned responses in courtship behavior of normal and mutant *Drosophila*. Proc Natl Acad Sci U S A 76, 3430–4 (1979).

39. Stocker, R. F. & Gendre, N. Courtship behavior of *Drosophila* genetically or surgically deprived of basiconic sensilla. Behav Genet. 19, 371–85 (1989).

40. McEwen, R. S. The reactions to light and to gravity in *Drosophila* and its mutants. J Exp Zool 25, 49–106 (1918).

41. Gailey, D. A., Jackson, F. R. & Siegel, R. W. Male courtship in *Drosophila*: the conditioned response to immature males and its genetic control. Genet. 102, 771–82 (1982).

42. McRobert, S. P. & Tompkins, L. Two consequences of homosexual courtship performed by *Drosophila melanogaster* and *Drosophila affinis males*. Evol. 42, 1093–7 (1988).

43. Zawistowski, S. & Richmond, R. C. Experience-mediated courtship reduction and competition for mates by male Drosophila melanogaster. Behav Genet. 15, 561–9 (1985).

44. Saleem, S., Ruggles, P. H., Abbott, W. K. & Carney, G. E. Sexual experience enhances *Drosophila melanogaster* male mating behavior and success. PLoS One 9, e96639 (2014). DOI 10.1371/journal.pone.0096639.

45. Manning, A. The control of sexual receptivity in female *Drosophila*. Anim Behav 15, 239–250 (1967).

46. Pan, Y. & Baker, B. S. Genetic identification and separation of innate and experience-dependent courtship behaviors in *Drosophila*. Cell 156, 236–248 (2014).

47. Liu, L., Davis, R. L. & Roman, G. Exploratory activity in *Drosophila* requires the *kurtz* nonvisual arrestin. Genet. 175, 1197–212 (2007). DOI 10.1534/genetics.106.068411.

48. Geer, B. & Green, M. Genotype, phenotype and mating behavior of *Drosophila melanogaster*. Am Nat 96, 175–181 (1962).

49. Ghosh, D. & Forrest, H. S. Enzymatic studies on the hydroxylation of kynurenine in *Drosophila melanogaster*. Genet. 55, 423–31 (1967).

50. Dreesen, T. D., Johnson, D. H. & Henikoff, S. The brown protein of *Drosophila melanogaster* is similar to the white protein and to components of active transport complexes. Mol Cell Biol 8, 5206–15 (1988).

51. Hersh, B. M. More than meets the eye: A primer for “Timing of locomotor recovery from anoxia modulated by the *white* gene in *Drosophila melanogaster*”. Genet. 204, 1369–1375 (2016).

52. Nolte, D. J. The eye-pigmentary system of *Drosophila*: The pigment cells. J Genet. 50, 79–99 (1950).

53. Pirrotta, V. & Bro¨ckl, C. Transcription of the *Drosophila white* locus and some of its mutants. EMBO J 3, 563–8 (1984).

54. Campbell, J. L. & Nash, H. A. Volatile general anesthetics reveal a neurobiological role for the *white* and *brown* genes of *Drosophila melanogaster*. J Neurobiol 49, 339–49 (2001).

55. Haddad, G. G., Sun, Y., Wyman, R. J. & Xu, T. Genetic basis of tolerance to O2 deprivation in *Drosophila melanogaster*. Proc Natl Acad Sci U S A 94, 10809–12 (1997).

56. Wang, H.-D., Kazemi-Esfarjani, P. & Benzer, S. Multiple-stress analysis for isolation of *Drosophila* longevity genes. Proc Natl Acad Sci U S A 101, 12610–12615 (2004).

57. Xiao, C., Mileva-Seitz, V., Seroude, L. & Robertson, R. M. Targeting hsp70 to motoneurons protects locomotor activity from hyperthermia in *Drosophila*. Dev Neurobiol 67, 438–455 (2007). DOI 10.1002/dneu.20344.

58. Zhang, S., Binari, R., Zhou, R. & Perrimon, N. A genomewide RNA interference screen for modifiers of aggregates formation by mutant Huntingtin in *Drosophila*. Genet. 184, 1165–79 (2010). DOI 10.1534/genetics.109.112516.

59. Hoenigsberg, H., Koref Santibanez, S. & Sironi, G. Intraspecific sexual preferences in *Drosophila prosaltans* Duda and in Drosophila equinoxialis Dobzhansky. Cell Mol Life Sci 15, 223–5 (1959).

60. Dukas, R. Male fruit flies learn to avoid interspecific courtship. Behav Ecol 15, 695–8 (2004).

